# Fast and memory efficient searching of large-scale mass spectrometry data using Tide

**DOI:** 10.1101/2025.04.01.646675

**Authors:** Attila Kertesz-Farkas, Frank Lawrence Nii Adoquaye Acquaye, Vladislav Ostapenko, Rufino Haroldo Locon, Yang Lu, Charles E. Grant, William Stafford Noble

**Affiliations:** Department of Data Analysis and Artificial Intelligence and Laboratory on AI for Computational Biology, Faculty of Computer Science, HSE University; University of Waterloo; Department of Genome Sciences, University of Washington; Paul G. Allen School of Computer Science and Engineering, University of Washington

**Keywords:** Tandem mass spectrometry, database search, peptide detection

## Abstract

Over the past 30 years, software for searching tandem mass spectrometry data against a protein database has improved dramatically in speed and statistical power. However, existing tools can still struggle to analyze truly massive datasets, when either the number of spectra or the number of proteins being analyzed grows too large. Here we describe enhancements to the Tide search engine that allow it to handle datasets containing >10 million spectra and databases containing >7 billion peptides on commodity hardware. We demonstrate that the new Tide architecture is around 2–7 times faster than the previous version and is now comparable to MSFragger and Sage in speed while requiring much less memory. Tide is open source and is publicly available as pre-compiled binaries for Windows, Linux and Mac.

## 1 Introduction

Database search is the killer app of proteomics tandem mass spectrometry analysis. Pioneered by the SEQUEST algorithm in 1994 [1], the core idea of iteratively scoring each observed MS2 spectrum against a database of peptides derived from a proteome database has been reimplemented in dozens of search tools that are used daily by proteomics mass spectrometry labs around the world (reviewed in [2]). Proteomics database search engines give rise to a host of related analytical questions, e.g., how to devise a score function that yields the highest quality results, how to recalibrate the PSM scores using additional sources of information, how to assess the statistical significance of the peptide-spectrum matches (PSMs) produced by the search engine, and how to aggregate those PSMs to the protein level.

In this work, we focus on a prior question: how can we make our search engine run efficiently, even when we need to analyze massive datasets? In particular, we address not just the speed of the search engine but its scalability. In general, the size of the input can be parameterized in two dimensions: the number of spectra that need to be searched and the number of peptides in the database. When either (or both) of these dimensions is very large, a good search engine should run efficiently and should not require too much memory. In practice, how much memory is “too much” depends on the available hardware. A few labs with access to large-scale computational infrastructure (or expertise and funding to work in a cloud-based setting) can work with computers with, say, 1 TB of random access memory (RAM). However, most commodity hardware has much less RAM.

Two popular and widely used search tools, MSFragger [3] and Sage [4], run very efficiently but require large amounts of RAM when analyzing large datasets. The exceptional speed offered by both tools comes from their use of the inverted indexing technique, pioneered by MSFragger, which is used to store and represent the theoretical fragment b- and y-ions of peptides. Unfortunately, this data structure has a memory footprint that scales approximately linearly in the size of the peptide database.

In this work, we focus on making a database search tool that works efficiently even with relatively limited memory consumption (say, up to 64 GB). To achieve this goal, we re-engineered the Tide search engine [5] from the ground up. In addition to simplifying and streamlining the code itself, the new implementation employs a distributed merge-and-sort operation on multiple input spectrum files, thereby limiting both the number of peptides and the number of spectra that must be kept in memory at once. In practice, we find that the new version of Tide is comparable in speed to MSFragger and Sage but is capable of analyzing datasets that are far too large to be analyzed using the inverted index data structure. Tide is also now capable of producing search results files in mzTab format [6], which includes support for specifying post-translational modifications (PTMs) using Unimod codes. The new version of Tide is available with an Apache open source license as part of the Crux Toolkit [7].

## 2 Methods

### 2.1 Code optimization

Since Tide’s initial publication in 2011 [5], the search code has been extended with various new features, such as the exact p-value calculation [8], tailor calibration [9], and residue-evidence scoring [10]. Each of these new features was implemented by a different developer. In the process, the code became fragmented, its complexity increased, and code maintenance became cumbersome. At the same time, the code modifications led to an overall decrease in Tide’s efficiency.

To address these problems, we restructured the Tide code and optimized the underlying data structures so that all components work smoothly together. In practice, Tide consists of two components: the tide-index command takes as input a protein fasta file and stores a corresponding peptide index on disk; the tide-search command then searches one or more spectrum files against the peptide index. The old tide-search program (version 4.2 and older) used an external class to generate the strings of modified and unmodified peptide sequences for reporting in the output results file. Furthermore, the code used a separate class to re-compute the theoretical fragmentation ions solely for the purpose of calculating the SEQUEST preliminary score (Sp). We eliminated these classes and the corresponding redundant calculations. Doing so halved the execution time in standard searches, i.e., when searching with a narrow precursor tolerance window using fully tryptic digestion.

During code optimization, we also aimed to reduce the number of user-level parameters in tide-search. The tailor calibration score and various statistics about matched b- and y-ions are now reported by default, eliminating the need for options (`--use-tailor-calibration` and `--compute-sp`) to turn these on and off. We also removed several parameters that are not widely used and of limited utility (`--file-column`, `--brief-output`, `--peptide-centric-search`, `--evidence-granularity`).

### 2.2 Optimization for speed and memory usage

Tide-search v4.2 (and older) was optimized for speed in searching one spectrum file at a time [5] (Figure 1A). The program takes as input a database of candidate peptides produced by tide-index. Tide-search iterates over the spectrum files, and in each iteration it first converts the experimental spectra into its own data format, sorts them by neutral mass, and stores them on disk. Finally, the program loops over the sorted spectra and scores them against their candidate peptides, which are read from disk from the index database. In this process, the candidate peptides are accessed using a rolling window technique, implemented with a double-ended queue container, which is adjusted for the spectrum’s neutral mass by discarding less massive peptides from the queue on one end and loading more massive peptides from the disk on the other end. The corresponding theoretical fragmentation ions are also calculated on the fly upon loading.

**Figure 1.**
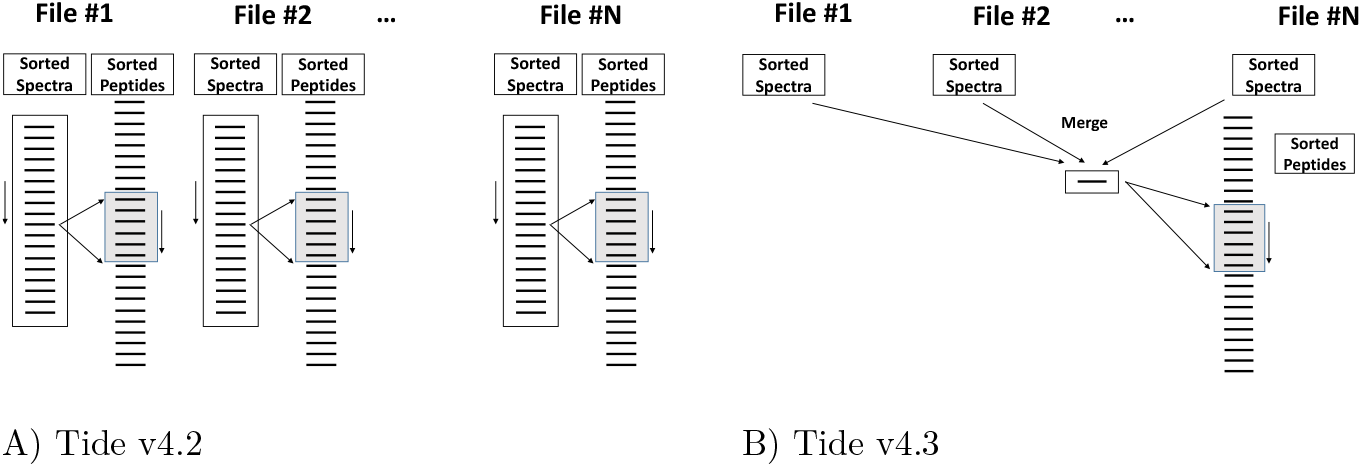
Architecture comparison of tide-search v4.2 and v4.3. **(A)** In the rolling window technique, the experimental spectra and the theoretical peptides are sorted by neutral mass in increasing order. Each experimental spectrum is iteratively searched against a window of theoretical peptides (shown as a grey box), which are determined by the precursor mass tolerance window parameter. The window of candidate peptides rolls over the peptide dataset by discarding less massive peptides from memory at one end and loading more massive peptides from disk at the other end of a double-ended queue container. Tide-search v4.2 iteratively processed the spectrum files, first sorting the spectra from one file in memory and then rolling over the peptides in the database, calculating theoretical fragmentation ions on the fly. Thus, each peptide’s fragment ions were recalculated for each spectrum file. **(B)** Tide-search v4.3 first sorts all spectra by neutral mass and stores the spectra on disk separately for each spectrum file. Spectra are then selected in sorted order from the preprocessed spectrum files. Scoring is done using the same rolling window technique. In the new approach, each peptide’s fragment ions are computed exactly once. This approach also results in a low memory footprint, because only one spectrum per file is kept in memory.

Unfortunately, tide-search v4.2 operates much less efficiently when analyzing multiple input spectrum files. The problem with its approach is two-fold. First, tide-search reads the entire peptide index data once for each spectrum file, reading from disk and recalculating the same theoretical fragmentation ions multiple times. Second, in settings such as metaproteomics or immunopeptidomics, the peptide index can occupy multiple terabytes on the disk [11], and the experimental spectrum data can be stored in hundreds of files. In such a setting, tide-search must stream terabytes of data from disk to memory for each spectrum file. For instance, searching experimental spectra stored in 100 mzML files against a terabyte peptide index database, tide-search v4.2 would stream 100 TB data from disk.

Accordingly, we have redesigned the architecture of tide-search avoid such redundant data streaming (Figure 1B). The new version of tide-search, v4.3, also converts the spectrum data to its own format, sorts the spectra by their neutral mass, and stores them on disk. However, instead of searching each file sequentially, tide-search v4.3 performs an external merge-and-sort operation on the fly to retrieve the spectra in sorted order from all the spectrum files. The consequence of this change is two-fold. First, tide-search will store only one spectrum in memory from each input file and stores them in a heap. Tide-search processes the least massive spectrum from the heap, and if it was retrieved from the *n*th input file, then it loads the next spectrum from the *n*th input file to the heap. Therefore, only *N* spectra are kept in memory for *N* input files, ensuring a low memory footprint. Second, the rolling window will go over the entire peptide index data only once during the whole search.

We note that both versions of tide-search perform similarly with respect to speed when provided with a single input spectrum file. We also note that both versions can use multi-threading during spectrum scoring.

### 2.3 Standardization of modifications and output format

In addition to improving tide-search’s efficiency, v4.3 is now capable of producing search results files in mzTab format [6]. This file format was designed by the Proteomics Standards Initiative (PSI) and aims to report results from a mass spectrometry proteomics experiment in a unified way. The format is designed to allow programmatic access to important data and metadata resulting from a given experiment. Tide-search’s mzTab output files have passed the validator at https://apps.lifs-tools.org/mztabvalidator/. This new format is activated using the `--mztab-output T` parameter. Tide-search still supports outputs in pepXML, mzIdentML, Percolator input (PIN), and its own tab-delimited format.

Part of the change to mzTab means that tide-search and tide-index now also allow modification specifications using Unimod codes (http://www.unimod.org/). For instance, to specify allowing up to one oxidation of methionine and one phosphorylation of serine, threonine or tyrosine, a user can specify `--mods-spec 1M[Unimod:35],1STY[Unimod:21]`. For backward compatibility, Tide still supports the old modification specification format, e.g., `--mods-spec 1M15.9949,1STY79.9663`, and these format types can be arbitrarily combined, e.g. `--mods-spec 1M15.9949,1STY[Unimod:21]`.

### 2.4 Data

To compare the performance of the various search engines, we use five publicly available data sets (Table 1). The first is a microbiome analysis (PXD011515 from PRIDE [18]), in which Rechenberger *et al*. analyzed 212 fecal samples from 56 hospitalized acute leukemia patients with multidrug-resistant Enterobactericeae (MRE) gut colonization [12]. In total, the dataset contains 12,475,565 spectra in 424 files. The protein database that Rechenberger *et al*. employed consisted of four parts: (a) the Integrated Genome Reference Catalog (`760MetaHit_139HMP_368PKU_511Bac.uniq.fasta`) with 9,878,647 entries, (b) the Swiss-Prot bacteria database (`uniprot-sprot_bac.fasta`) with 333,480 entries, (c) the Swiss-Prot human canonical sequences database (`uniprot_cano_varsplic_HUMAN.fasta`) 43,123 entries, and (d) 20,720,510 metaproteomic protein sequences from 189 fasta files (with names containing `ORF`). The fasta files are available at the PXD011515 project at PRIDE. The concatenation of these fasta files results in 30,975,760 proteins, and *in silico* digestion allowing oxidation and acetylation as variable modifications resulted in 7,104,692,523 distinct peptides.

**Table 1:** Data sets. “Q-Ex” stands for Q-Exactive. PTMs correspond to oxidation of methionine (Ox), phosphorylation of serine, threonine and tyrosine (Ph), lysine and N-terminal acetylation (Ac), deamidation of asparagine and glutamine (Dea), c-terminal amidation (Am) and TMT-6plex tags (TMT). The column “Files (GB)” indicates the number of spectrum files mzML format, and their total size in GB. The column “Peptides (GB)” indicates the number of peptides generated by tide-index, and the size of its data on disk. “APC” is the average number of candidate peptides as reported by tide-search. The three accessions listed in lines 2–5 of the table jointly comprise the PEAKS data set.

| Accession | Database | Instrument | PTMs | Spectra | Files (GB) | Peptides (GB) | ACP |
| --- | --- | --- | --- | --- | --- | --- | --- |
| PXD011515 [12] | Metaproteome | Q-Ex. HF-X | Ox, Ac | 12,475,565 | 424 (445) | 7,104,692,523 (461) | 272,257 |
| MSV000084172 [13] | *H. sapiens* | Q-Ex. Plus | Ox, Dea, Ac | 16,903,733 | 371 (198) | 5,796,938,142 (410) | 17,674 |
| MSV000080527 [13] |  |  |  | 3,757,498 | 87 (30) |  |  |
| PXD004894 [14] |  |  |  | 5,294,250 | 133 (70) |  |  |
| PXD017407 [15] | *H. sapiens* | Q Ex. HF-X | Ox, Ph, Ac, TMT, Am | 673,764 | 21 (5) | 2,353,638,986 (128) | 133,118 |
| PXD019483 [16] | *H. sapiens* | Q Ex. HF-X | Ox, Ph, Ac, | 22,473,687 | 127 (1359) | 142,245,554 (8.0) | 2,511 |
| PXD028806 [17] | *H. sapiens* | Q-Ex. Plus | Ox, TMT | 1,410,179 | 24 (221) | 11,519,178 (0.7) | 270 |

The second case study was created to demonstrate the scalability of the PEAKS analysis software [19]. This study was constructed from three immunopeptidomics experiments (MSV000084172, MSV000080527, PXD004894), consisting of a total of 25,249,574 spectra stored in 539 spectrum files. The data were downloaded from the PRIDE and MassIVE databases. The human proteme database, `uniprot-proteome_ UP000005640_canonical_isoforms.fasta` containing 101,038 proteins, was downloaded from Uniprot [20] on March 22, 2022. After non-specific *in silico* digestion, 5,796,938,142 non-tryptic peptides of length between 7 and 40 were obtained, allowing oxidation of methonine, acetylation of lysine and N-terminal residues, and deamidation of asparagine and glutamine (Dea) as variable modifications.

In addition, we employed three smaller datasets from the PRIDE repository. The PXD017407 dataset [15] was derived from an analysis of peptides bound to class I major histocompatibility complexes (MHC). This dataset contains 705,218 spectra with high resolution precursor and fragmentation information in 21 mzML files. Peptides were labeled with TMT6plex and analyzed using a Thermo Fisher Q Exactive HF-X Hybrid Quadrupole-Orbitrap mass spectrometer in DDA mode. For this and the subsequent two datasets, we used the `uniprot-proteome UP000005640.fasta` file downloaded from Uniprot [20] on March 22, 2022. This fasta file contains 79,071 canonical protein sequences. The corresponding peptide index data is generated with non-typtic cleavage, allowing peptides of length 7–15 amino acids and including oxidized methionine and phosphorylated serine, threonine, and tyrosine as variable modifications. The resulting index contains 2,353,638,986 peptides, which occupied 128 GB of disk space.

The PXD019483 dataset [16] was obtained from a proteome identification and quantification study, in which the peptide separation step was performed by a microstructured and extremely reproducible chromatographic system, for the in-depth measurement of 100 taxonomically diverse organisms. We focused solely on the human data. The peptide index data was generated from the human protein sequences via tryptic digestion, allowing peptides of length 7–50 amino acids and including oxidized methionine, phosphorylated serine, threonine, and tyrosine; acetylation of peptide N-terminal residues and lysine amino acids. The resulting index contains 142,245,554 peptides, which took 8 GB of disk space.

The PXD028806 dataset [17] was obtained from two breast cancer cell lines with different levels of glycolytic activity. Peptides were labeled with TMT tags and analyzed with a Q Exactive Plus instrument. The spectra were stored in 24 files, occupying 221 GB of disk space. The corresponding peptide index data contained 11,519,178 peptides with lengths 7–50 amino acids, obtained with tryptic cleavage, allowing up to 3 missed cleavages, and including oxidixed methionine as variable modification.

For each data set, raw files were converted to mzML format with using the msconvert program from ProteoWizard toolkit [21]. Search parameter settings were taken from the associated papers, as summarized in Table 1. Carbamidomethylation of cysteine was specified as a fixed modification for all searches. For all datasets, the precursor tolerance window and fragment ion tolerance windows were set 10 pmm and 0.02 Da, respectively.

### 2.5 Database search

All database searches were performed using the tide-search command (v4.2 and v4.3), MSFragger v3.7, and Sage v0.14.4. The output files from tide-search and MSFragger were analyzed by Percolator version 3.06.01 [22] embedded in the Crux toolkit v4.2. Sage employed its own built-in PSM rescoring method, based on linear discriminant analysis, that is similar to Percolator. All methods were executed with 4 CPU threads. Log files for tide-index and tide-search containing the full command specification and parameterization, as well as Sage and MSFragger parameter files, are available in the zipped supplementary files.

All our experiments were executed on a Linux server equipped with an Intel Xeon CPU E5-2640 v4 2.40GHz processor with 20 cores and 128 GB DDR RAM, operated by Ubuntu 22.04.4 LTS.

### 2.6 False discovery rate control

False discovery rate (FDR) control was performed using target-decoy competition. For Tide, the decoy peptide sequences were generated automatically by tide-index by reversing the non-terminal amino acids of the target peptides. Occasionally, the target database contains two peptides that are reversed versions of one another; in such a setting, the corresponding decoys are created by shuffling rather than reversal. MS-Fragger does not generate decoy peptides internally; therefore, we concatenated the reversed protein sequences as decoys to the corresponding fasta files. Sage generates its own decoy peptides by reversing the internal amino acids of the target peptides.

The FDR control was performed by Percolator in the case of tide-search and MS-Fragger, and by Sage. We have recently demonstrated that PSM-level FDR control is problematic due to dependencies between observed spectra [23]; therefore, we report the number of peptides accepted at 1% peptide-level FDR, reported by either Percolator or Sage.

## 3 Results

### 3.1 Microbiome data analysis

We began by using Tide to repeat a previously reported microbiome analysis, in which Rechenberger *et al*. analyzed 212 fecal samples from 56 hospitalized acute leukemia patients with multidrug-resistant *Enterobactericeae* (MRE) gut colonization [12] (PXD011515). In total, the dataset contains 12.5 million spectra, and the corresponding database contains 31 million proteins. Not surprisingly, searching this massive set of spectra against this massive database is expensive. Rechenberger *et al*. used MaxQuant [24] for their analysis, and they report that the searches required 1834 h (10 weeks) of run time using 10 cores on a Intel Xeon Gold 6150 CPU at 2.70GHz with 4 processors. We repeated this analysis using our server with similar search parameters, and the analysis required 179 hours (7.5 days) for Tide v4.3. Taking into account that we used 4 cores rather than 10, this still represents more than a 5-fold speedup relative to MaxQuant. This analysis required 1,133 hours (6.7 weeks) for Tide v4.2, representing a >6-fold speedup.

Notably, we also attempted to run Sage and MSFragger on this dataset, but neither could handle the very large protein database, even after we switched to a system with 1536 GB of RAM [25] (Supplementary files: `logs/PXD011515/resource_usage_MSFragger.jpg` and `logs/PXD011515/resource_usage_ SAGE.jpg`) MSFragger does not allow generating a database with more than two billion peptides, so we manually split the fasta file into 32 batches, executed MSFragger with each batch sequentially, and then merged the search results by keeping the spectrum annotations that provided the smallest `log10 evalue` from the batch searches. MSFragger used 124 GB of RAM to analyze a single batch, and it required a total of 448 hours of execution time for the whole dataset. We note that MSFragger executed the actual spectrum search fast; however, it took significantly more time to carry out remapping alternative proteins and postprocessing. The new version of tide-search analyzed this dataset in a single step, and it required slightly less than 20 GB of RAM and 179 hours, representing a >2-fold speedup. We note that MSFragger ran out of memory when we split the fasta files into a smaller number of batches. Tide-search v4.2 and v4.3 programs resulted in around 351 thousands peptide identifications at 1% FDR, while MSFragger resulted in only 210 thousand peptides (Table 2). We speculate that MSFragger may have performed less well because the E-value calculation is based on the histogram of scores produced by each spectrum. Dividing the database into separate subsets yields 32 smaller histograms and, hence, variably calibrated E-values for a single spectrum. We note that the FragPipe GUI offers a `split-dataset` option to run searches against large peptide datasets; however, at the time we ran our experiments, this option failed to run successfully due. Sage failed with an out-of-memory error even when searching one of the 32 batches.

**Table 2:** Comparison of search engines. The table lists the execution time in seconds, memory usage in gigabytes, and the number of detected peptides at 1% peptide-level FDR. “PEAKS data” refers to the concatenation of MSV000084172, PXD004894, PXD022950. Missing tests (—) failed due to out of memory issues. The amount of available RAM was 128 GB. ^1^For MSFragger, the protein fasta file was split into 32 batches for the PXD011515, and into 2 batches for the PXD017407. Results indicate total execution time of all batches and memory usage for processing a single batch. ^2^Estimate. We aborted this test due to CPU time limit. Estimation is based on 278 spectrum files (out of 571) being searched in 25 days. Peptide detection was estimated based on Tide-search v4.3. results.

|  | Tide-search v4.3 |  |  | Tide-search v4.2 |  |  | Sage |  |  | MSFragger<sup>1</sup> |  |  |
| --- | --- | --- | --- | --- | --- | --- | --- | --- | --- | --- | --- | --- |
|  | Time | RAM | Peptides | Time | RAM | Peptides | Time | RAM | Peptides | Time | RAM | Peptides |
| PXD011515 | 645,000 | 17 | 358,670 | 4,080,000 | 16 | 351,118 | — | — | — | 1,612,620 | 124 | 210,056 |
| PEAKS data | 26,600 | 2 | 173,712 | 4,436,546<sup>2</sup> | 1 | 173,712<sup>2</sup> | — | — | — | — | — | — |
| PXD017407 | 18,700 | 3 | 15,137 | 60,900 | 3 | 15,045 | — | — | — | 3,360 | 98 | 15,958 |
| PXD019483 | 12,900 | 9 | 170,519 | 42,500 | 2 | 170,645 | — | — | — | 14,700 | 84 | 197,810 |
| PXD028806 | 673 | 9 | 114,158 | 1420 | 2 | 114,205 | 431 | 15 | 113,321 | 793 | 64 | 119,309 |

### 3.2 Immunopeptidomics data analysis

The second case study was created to demonstrate the scalability of the PEAKS analysis software [19]. This study was constructed from three experiments (MSV000084172, MSV000080527, PXD004894), consisting of a total of 25.2 million spectra stored in 539 spectrum files. The human proteome database that this study used contained 101,038 proteins and, after non-specific *in silico* digestion, 5.8 billion non-tryptic peptides. In the initial study, the search was run on the Amazon Web Services cloud with 512 cores and 1 TB RAM in 17 hours, not including the time required to upload the data. Tide required 8.6 hours to generate the peptide index data and 7.4 hours to search the data using 4 threads and requiring only 2 GB of RAM; a total of 16.3 hours. Thus, Tide is comparable to PEAKS in speed, despite using only 4 threads rather than 512 and 2 GB rather than 1 TB of RAM. Notably, the old version of tide-search required 25 days to search half of the spectrum data (278 spectrum files out of the 571 files–we halted this search due to limited CPU time availability). Therefore, for this dataset tide-search v4.3 is ∼ 160 times faster than v4.2 (1232 vs. 7.2 hours) (Table 2). We also attempted to run Sage and MSFragger on this dataset, but neither could handle the very large peptide database. We note that we tried to ensure the same parameterization of tide-search and PEAKS, but some parameters (e.g., allowed peptide lengths and missed cleavages, maximum number of variable modifications per peptide) were not specified in the original study.

### 3.2 Human proteome analysis

Finally, we searched the three, smaller datasets from PRIDE (PXD017407, PXD019483, PXD028806) to carry out a direct comparison of MSFragger, Sage and both versions of Tide on the same hardware (Table 2). Sage successfully ran to completion only with the PXD028806 dataset, running out of memory for the other two datasets. On this dataset, Sage was somewhat faster than tide-search v4.3 (431 s. versus 673 s.), though Sage required more RAM (15 GB versus 9 GB). On the same dataset, MSFragger was slower than either of the other two tools (793 s.) and required substantially more RAM (64 GB). The new tide-search was also slightly faster than MSFragger on the PXD019483 dataset. The MSFragger was >5-fold faster than tide-search on PXD017407 (3,360 s. versus 18,600 s.) because the inverted fragmentation indexing makes the scoring of one spectrum against a very large number of candidate peptides computationally efficient, which is the case with this dataset. In general, Tide’s RAM consumption is around 10 time less than what is required for MSFragger, and much less than Sage’s consumption.

Comparing the old versus new versions of tide-search, we observe that the new version (v4.3) requires more RAM than the old version (v4.2). This is because the spectrum file parsing now requires keeping the whole file in memory when sorting the experimental spectra by neutral mass. However, during the actual spectrum scoring against the peptide database, tide-search v4.3 uses only 1–2 GB. The timing results suggest that the new version of tide-search is approximately 2–7 times faster than the previous version.

We also note that MSFragger tends to yield 5–10% more peptide detections at 1% FDR level than tide-search. This improvement is likely due to the additional PSM features that MSFragger provides as input to Percolator, thought it may also relate to the use of a different score function (hyperscore versus XCorr) and a different score calibration procedure (E-value versus tailor calibration).

Finally, we mention that in these experiments Tide used a conventional hard drive to store the spectra and the peptide indices. Tide would likely run substantially faster if the data could instead be stored on a solid state drive.

## 4 Discussion

Currently popular database search tools, such as MSFragger, Sage, MaxQuant, and Comet [26], are fast enough to perform standard database searching procedures, i.e., with a narrow tolerance window, with tryptic peptides harboring a few PTMs, and with a relatively small protein database. However, these tools become difficult or impossible to use with very large datasets. We have scaled up the Tide search engine to efficiently handle datasets that are large both in terms of the number of spectra and the number of peptides in the index.

Over a year ago, we updated the peptide database indexing step in tide-index [11], so that it can now efficiently generate very large peptide datasets. In that work, we demonstrated that the new version tide-index requires around 3–5 times less CPU time and memory than the previous version, and it can now generate very large peptide databases, subject only to the availability of disk space. In parallel with this project we also overhauled the tide-search code with a focus on speeding up its scoring functions up, yielding a three-fold increase in speed in scoring relative to the previous version of tide-search [27]. Now in this current project, we have redesigned the whole architecture of tide-search so that it can efficiently search tera-scale MS2 data against tera-scale peptide indices.

We conclude that the exceptional speed of MSFragger and Sage comes from the use of the inverted fragment ion index to represent the theoretical fragment b- and y-ions of peptides. This data structure is efficient but requires that the whole peptide index is stored in memory throughout the search. Thus, MS-fragger and Sage achieve their exceptional speed at the expense of memory consumption. Tide-search is also on par with Sage and MSFragger in terms of speed while requiring much less memory due to its rolling window scoring procedure. We demonstrated in our benchmark test that the new Tide architecture is around 2–7 times faster than the previous version and is now comparable to MSFragger and Sage in speed while requiring much less memory. Based on published timing results on our benchmark data, we also estimate that Tide is ∼ 10–20 times faster than MaxQuant [24].

We have also modified Tide to adopt several mass spectrometry standards. The search engine can now report results in mzTab format, and it also can handle modifications with Unimod identifiers. We aim to continue actively supporting Tide and facilitating its adoption by the mass spectrometry community. We are also actively working on integrating Tide into several widely used software tools, including Skyline [28], SearchGUI [29], PeptideShaker [30], and quantms [31], which will allow users to seamlessly integrate Tide into their workflows.

## Supporting information

Supplementary log files

## Acknowledgment

This research was supported in part through computational resources provided by the HPC facilities at HSE University [25] and by National Science Foundation award 2245300.

## Author contributions

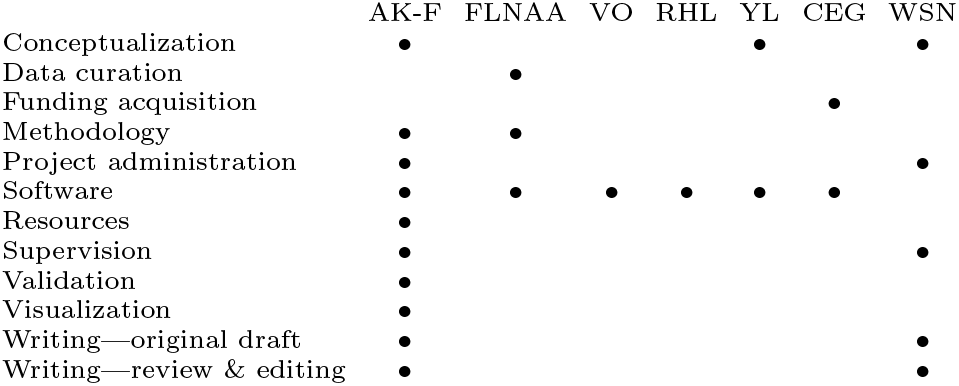

## Availability

Tide-search is available in the latest version of the Crux toolkit at https://crux.ms. Apache licensed source code is available, and pre-compiled binaries are provided for Windows, MacOS and Linux, accessible via the linked github repository (https://github.com/crux-toolkit/crux-toolkit/).

## Supporting information

The following supporting information is available free of charge at ACS website http://pubs.acs.org

1. Supplementary Files: Zipped log files of tide-search and tide-index that also contains the fully parameterized and specified commands, as well as Sage and MS-Fragger parameter files.

