## Supplementary log files for "Fast and memory efficient searching of large-scale mass spectrometry data using Tide": tide-index.log.txt

INFO: CPU: L429746
INFO: Crux version: 4.1-b64e8589-2024-02-02
INFO: Вс 04 фев 2024 02:19:18 MSK
COMMAND LINE: /home/attila/crux-tide-lite/src/crux tide-index --max-mods 2 --mods-spec 1M+15.9949,1K+229.162932,1K+42.010565,1STY+79.966331 --nterm-peptide-mods-spec X+229.162932 --cterm-peptide-mods-spec 1X-0.98402 --decoy-format shuffle --digestion non-specific-digest --missed-cleavages 0 --min-length 7 --max-length 15 --overwrite T --peptide-list F --temp-dir /hdd/data/tide-lite-results/HUMAN_NT_1-15_OxPhosAcetyl_deamid_TMT_TL --output-dir /hdd/data/tide-lite-results/HUMAN_NT_1-15_OxPhosAcetyl_deamid_TMT_TL /home/data/Fasta/uniprot-proteome_UP000005640.fasta /hdd/data/tide-lite-results/HUMAN_NT_1-15_OxPhosAcetyl_deamid_TMT_TL
INFO: Beginning tide-index.
INFO: Running tide-index...
WARNING: The output directory '/hdd/data/tide-lite-results/HUMAN_NT_1-15_OxPhosAcetyl_deamid_TMT_TL' already exists.
Existing files will be overwritten.
INFO: Reading /home/data/Fasta/uniprot-proteome_UP000005640.fasta and computing unmodified target peptides...
INFO: Processed 10000 protein sequences
INFO: Processed 20000 protein sequences
INFO: Processed 30000 protein sequences
INFO: Processed 40000 protein sequences
INFO: Processed 50000 protein sequences
INFO: Processed 60000 protein sequences
INFO: Processed 70000 protein sequences
INFO: Cleaved 79052 protein sequences in total.
INFO: Ignoring 77475 peptide sequences containing unrecognized characters.
INFO: Generated 244814916 targets, including duplicates.
INFO: Writing peptides
INFO: Wrote 1000000 unique target peptides
INFO: Wrote 2000000 unique target peptides
INFO: Wrote 3000000 unique target peptides
INFO: Wrote 4000000 unique target peptides
INFO: Wrote 5000000 unique target peptides
INFO: Wrote 6000000 unique target peptides
INFO: Wrote 7000000 unique target peptides
INFO: Wrote 8000000 unique target peptides
INFO: Wrote 9000000 unique target peptides
INFO: Wrote 10000000 unique target peptides
INFO: Wrote 11000000 unique target peptides
INFO: Wrote 12000000 unique target peptides
INFO: Wrote 13000000 unique target peptides
INFO: Wrote 14000000 unique target peptides
INFO: Wrote 15000000 unique target peptides
INFO: Wrote 16000000 unique target peptides
INFO: Wrote 17000000 unique target peptides
INFO: Wrote 18000000 unique target peptides
INFO: Wrote 19000000 unique target peptides
INFO: Wrote 20000000 unique target peptides
INFO: Wrote 21000000 unique target peptides
INFO: Wrote 22000000 unique target peptides
INFO: Wrote 23000000 unique target peptides
INFO: Wrote 24000000 unique target peptides
INFO: Wrote 25000000 unique target peptides
INFO: Wrote 26000000 unique target peptides
INFO: Wrote 27000000 unique target peptides
INFO: Wrote 28000000 unique target peptides
INFO: Wrote 29000000 unique target peptides
INFO: Wrote 30000000 unique target peptides
INFO: Wrote 31000000 unique target peptides
INFO: Wrote 32000000 unique target peptides
INFO: Wrote 33000000 unique target peptides
INFO: Wrote 34000000 unique target peptides
INFO: Wrote 35000000 unique target peptides
INFO: Wrote 36000000 unique target peptides
INFO: Wrote 37000000 unique target peptides
INFO: Wrote 38000000 unique target peptides
INFO: Wrote 39000000 unique target peptides
INFO: Wrote 40000000 unique target peptides
INFO: Wrote 41000000 unique target peptides
INFO: Wrote 42000000 unique target peptides
INFO: Wrote 43000000 unique target peptides
INFO: Wrote 44000000 unique target peptides
INFO: Wrote 45000000 unique target peptides
INFO: Wrote 46000000 unique target peptides
INFO: Wrote 47000000 unique target peptides
INFO: Wrote 48000000 unique target peptides
INFO: Wrote 49000000 unique target peptides
INFO: Wrote 50000000 unique target peptides
INFO: Wrote 51000000 unique target peptides
INFO: Wrote 52000000 unique target peptides
INFO: Wrote 53000000 unique target peptides
INFO: Wrote 54000000 unique target peptides
INFO: Wrote 55000000 unique target peptides
INFO: Wrote 56000000 unique target peptides
INFO: Wrote 57000000 unique target peptides
INFO: Wrote 58000000 unique target peptides
INFO: Wrote 59000000 unique target peptides
INFO: Wrote 60000000 unique target peptides
INFO: Wrote 61000000 unique target peptides
INFO: Wrote 62000000 unique target peptides
INFO: Wrote 63000000 unique target peptides
INFO: Wrote 64000000 unique target peptides
INFO: Wrote 65000000 unique target peptides
INFO: Wrote 66000000 unique target peptides
INFO: Wrote 67000000 unique target peptides
INFO: Wrote 68000000 unique target peptides
INFO: Wrote 69000000 unique target peptides
INFO: Wrote 70000000 unique target peptides
INFO: Wrote 71000000 unique target peptides
INFO: Wrote 72000000 unique target peptides
INFO: Wrote 73000000 unique target peptides
INFO: Wrote 74000000 unique target peptides
INFO: Wrote 75000000 unique target peptides
INFO: Wrote 76000000 unique target peptides
INFO: Wrote 77000000 unique target peptides
INFO: Wrote 78000000 unique target peptides
INFO: Wrote 79000000 unique target peptides
INFO: Wrote 80000000 unique target peptides
INFO: Wrote 81000000 unique target peptides
INFO: Wrote 82000000 unique target peptides
INFO: Wrote 83000000 unique target peptides
INFO: Wrote 84000000 unique target peptides
INFO: Wrote 85000000 unique target peptides
INFO: Wrote 86000000 unique target peptides
INFO: Wrote 87000000 unique target peptides
INFO: Wrote 88000000 unique target peptides
INFO: Wrote 89000000 unique target peptides
INFO: Wrote 90000000 unique target peptides
INFO: Wrote 91000000 unique target peptides
INFO: Wrote 92000000 unique target peptides
INFO: Wrote 93000000 unique target peptides
INFO: Wrote 94000000 unique target peptides
INFO: Wrote 95000000 unique target peptides
INFO: Wrote 96000000 unique target peptides
INFO: Wrote 97000000 unique target peptides
INFO: Wrote 98000000 unique target peptides
INFO: Wrote 99000000 unique target peptides
INFO: Wrote 100000000 unique target peptides
INFO: Wrote 101000000 unique target peptides
INFO: Wrote 102000000 unique target peptides
INFO: Skipped 142099339 duplicate targets.
INFO: Generated 102715577 unique target peptides.
INFO: Computing modified peptides...
INFO: Wrote 10000000 modified target peptides to temp files
INFO: Wrote 20000000 modified target peptides to temp files
INFO: Wrote 30000000 modified target peptides to temp files
INFO: Wrote 40000000 modified target peptides to temp files
INFO: Wrote 50000000 modified target peptides to temp files
INFO: Wrote 60000000 modified target peptides to temp files
INFO: Wrote 70000000 modified target peptides to temp files
INFO: Wrote 80000000 modified target peptides to temp files
INFO: Wrote 90000000 modified target peptides to temp files
INFO: Wrote 100000000 modified target peptides to temp files
INFO: Wrote 110000000 modified target peptides to temp files
INFO: Wrote 120000000 modified target peptides to temp files
INFO: Wrote 130000000 modified target peptides to temp files
INFO: Wrote 140000000 modified target peptides to temp files
INFO: Wrote 150000000 modified target peptides to temp files
INFO: Wrote 160000000 modified target peptides to temp files
INFO: Wrote 170000000 modified target peptides to temp files
INFO: Wrote 180000000 modified target peptides to temp files
INFO: Wrote 190000000 modified target peptides to temp files
INFO: Wrote 200000000 modified target peptides to temp files
INFO: Wrote 210000000 modified target peptides to temp files
INFO: Wrote 220000000 modified target peptides to temp files
INFO: Wrote 230000000 modified target peptides to temp files
INFO: Wrote 240000000 modified target peptides to temp files
INFO: Wrote 250000000 modified target peptides to temp files
INFO: Wrote 260000000 modified target peptides to temp files
INFO: Wrote 270000000 modified target peptides to temp files
INFO: Wrote 280000000 modified target peptides to temp files
INFO: Wrote 290000000 modified target peptides to temp files
INFO: Wrote 300000000 modified target peptides to temp files
INFO: Wrote 310000000 modified target peptides to temp files
INFO: Wrote 320000000 modified target peptides to temp files
INFO: Wrote 330000000 modified target peptides to temp files
INFO: Wrote 340000000 modified target peptides to temp files
INFO: Wrote 350000000 modified target peptides to temp files
INFO: Wrote 360000000 modified target peptides to temp files
INFO: Wrote 370000000 modified target peptides to temp files
INFO: Wrote 380000000 modified target peptides to temp files
INFO: Wrote 390000000 modified target peptides to temp files
INFO: Wrote 400000000 modified target peptides to temp files
INFO: Wrote 410000000 modified target peptides to temp files
INFO: Wrote 420000000 modified target peptides to temp files
INFO: Wrote 430000000 modified target peptides to temp files
INFO: Wrote 440000000 modified target peptides to temp files
INFO: Wrote 450000000 modified target peptides to temp files
INFO: Wrote 460000000 modified target peptides to temp files
INFO: Wrote 470000000 modified target peptides to temp files
INFO: Wrote 480000000 modified target peptides to temp files
INFO: Wrote 490000000 modified target peptides to temp files
INFO: Wrote 500000000 modified target peptides to temp files
INFO: Wrote 510000000 modified target peptides to temp files
INFO: Wrote 520000000 modified target peptides to temp files
INFO: Wrote 530000000 modified target peptides to temp files
INFO: Wrote 540000000 modified target peptides to temp files
INFO: Wrote 550000000 modified target peptides to temp files
INFO: Wrote 560000000 modified target peptides to temp files
INFO: Wrote 570000000 modified target peptides to temp files
INFO: Wrote 580000000 modified target peptides to temp files
INFO: Wrote 590000000 modified target peptides to temp files
INFO: Wrote 600000000 modified target peptides to temp files
INFO: Wrote 610000000 modified target peptides to temp files
INFO: Wrote 620000000 modified target peptides to temp files
INFO: Wrote 630000000 modified target peptides to temp files
INFO: Wrote 640000000 modified target peptides to temp files
INFO: Wrote 650000000 modified target peptides to temp files
INFO: Wrote 660000000 modified target peptides to temp files
INFO: Wrote 670000000 modified target peptides to temp files
INFO: Wrote 680000000 modified target peptides to temp files
INFO: Wrote 690000000 modified target peptides to temp files
INFO: Wrote 700000000 modified target peptides to temp files
INFO: Wrote 710000000 modified target peptides to temp files
INFO: Wrote 720000000 modified target peptides to temp files
INFO: Wrote 730000000 modified target peptides to temp files
INFO: Wrote 740000000 modified target peptides to temp files
INFO: Wrote 750000000 modified target peptides to temp files
INFO: Wrote 760000000 modified target peptides to temp files
INFO: Wrote 770000000 modified target peptides to temp files
INFO: Wrote 780000000 modified target peptides to temp files
INFO: Wrote 790000000 modified target peptides to temp files
INFO: Wrote 800000000 modified target peptides to temp files
INFO: Wrote 810000000 modified target peptides to temp files
INFO: Wrote 820000000 modified target peptides to temp files
INFO: Wrote 830000000 modified target peptides to temp files
INFO: Wrote 840000000 modified target peptides to temp files
INFO: Wrote 850000000 modified target peptides to temp files
INFO: Wrote 860000000 modified target peptides to temp files
INFO: Wrote 870000000 modified target peptides to temp files
INFO: Wrote 880000000 modified target peptides to temp files
INFO: Wrote 890000000 modified target peptides to temp files
INFO: Wrote 900000000 modified target peptides to temp files
INFO: Wrote 910000000 modified target peptides to temp files
INFO: Wrote 920000000 modified target peptides to temp files
INFO: Wrote 930000000 modified target peptides to temp files
INFO: Wrote 940000000 modified target peptides to temp files
INFO: Wrote 950000000 modified target peptides to temp files
INFO: Wrote 960000000 modified target peptides to temp files
INFO: Wrote 970000000 modified target peptides to temp files
INFO: Wrote 980000000 modified target peptides to temp files
INFO: Wrote 990000000 modified target peptides to temp files
INFO: Wrote 1000000000 modified target peptides to temp files
INFO: Wrote 1010000000 modified target peptides to temp files
INFO: Wrote 1020000000 modified target peptides to temp files
INFO: Wrote 1030000000 modified target peptides to temp files
INFO: Wrote 1040000000 modified target peptides to temp files
INFO: Wrote 1050000000 modified target peptides to temp files
INFO: Wrote 1060000000 modified target peptides to temp files
INFO: Wrote 1070000000 modified target peptides to temp files
INFO: Wrote 1080000000 modified target peptides to temp files
INFO: Wrote 1090000000 modified target peptides to temp files
INFO: Wrote 1100000000 modified target peptides to temp files
INFO: Wrote 1110000000 modified target peptides to temp files
INFO: Wrote 1120000000 modified target peptides to temp files
INFO: Wrote 1130000000 modified target peptides to temp files
INFO: Wrote 1140000000 modified target peptides to temp files
INFO: Wrote 1150000000 modified target peptides to temp files
INFO: Wrote 1160000000 modified target peptides to temp files
INFO: Wrote 1170000000 modified target peptides to temp files
INFO: Created 1177005847 modified and unmodified target peptides.
INFO: Generating 1 decoy(s) per target peptide
INFO: Wrote 10000000 target and their corresponding decoy peptides
INFO: Wrote 20000000 target and their corresponding decoy peptides
INFO: Wrote 30000000 target and their corresponding decoy peptides
INFO: Wrote 40000000 target and their corresponding decoy peptides
INFO: Wrote 50000000 target and their corresponding decoy peptides
INFO: Wrote 60000000 target and their corresponding decoy peptides
INFO: Wrote 70000000 target and their corresponding decoy peptides
INFO: Wrote 80000000 target and their corresponding decoy peptides
INFO: Wrote 90000000 target and their corresponding decoy peptides
INFO: Wrote 100000000 target and their corresponding decoy peptides
INFO: Wrote 110000000 target and their corresponding decoy peptides
INFO: Wrote 120000000 target and their corresponding decoy peptides
INFO: Wrote 130000000 target and their corresponding decoy peptides
INFO: Wrote 140000000 target and their corresponding decoy peptides
INFO: Wrote 150000000 target and their corresponding decoy peptides
INFO: Wrote 160000000 target and their corresponding decoy peptides
INFO: Wrote 170000000 target and their corresponding decoy peptides
INFO: Wrote 180000000 target and their corresponding decoy peptides
INFO: Wrote 190000000 target and their corresponding decoy peptides
INFO: Wrote 200000000 target and their corresponding decoy peptides
INFO: Wrote 210000000 target and their corresponding decoy peptides
INFO: Wrote 220000000 target and their corresponding decoy peptides
INFO: Wrote 230000000 target and their corresponding decoy peptides
INFO: Wrote 240000000 target and their corresponding decoy peptides
INFO: Wrote 250000000 target and their corresponding decoy peptides
INFO: Wrote 260000000 target and their corresponding decoy peptides
INFO: Wrote 270000000 target and their corresponding decoy peptides
INFO: Wrote 280000000 target and their corresponding decoy peptides
INFO: Wrote 290000000 target and their corresponding decoy peptides
INFO: Wrote 300000000 target and their corresponding decoy peptides
INFO: Wrote 310000000 target and their corresponding decoy peptides
INFO: Wrote 320000000 target and their corresponding decoy peptides
INFO: Wrote 330000000 target and their corresponding decoy peptides
INFO: Wrote 340000000 target and their corresponding decoy peptides
INFO: Wrote 350000000 target and their corresponding decoy peptides
INFO: Wrote 360000000 target and their corresponding decoy peptides
INFO: Wrote 370000000 target and their corresponding decoy peptides
INFO: Wrote 380000000 target and their corresponding decoy peptides
INFO: Wrote 390000000 target and their corresponding decoy peptides
INFO: Wrote 400000000 target and their corresponding decoy peptides
INFO: Wrote 410000000 target and their corresponding decoy peptides
INFO: Wrote 420000000 target and their corresponding decoy peptides
INFO: Wrote 430000000 target and their corresponding decoy peptides
INFO: Wrote 440000000 target and their corresponding decoy peptides
INFO: Wrote 450000000 target and their corresponding decoy peptides
INFO: Wrote 460000000 target and their corresponding decoy peptides
INFO: Wrote 470000000 target and their corresponding decoy peptides
INFO: Wrote 480000000 target and their corresponding decoy peptides
INFO: Wrote 490000000 target and their corresponding decoy peptides
INFO: Wrote 500000000 target and their corresponding decoy peptides
INFO: Wrote 510000000 target and their corresponding decoy peptides
INFO: Wrote 520000000 target and their corresponding decoy peptides
INFO: Wrote 530000000 target and their corresponding decoy peptides
INFO: Wrote 540000000 target and their corresponding decoy peptides
INFO: Wrote 550000000 target and their corresponding decoy peptides
INFO: Wrote 560000000 target and their corresponding decoy peptides
INFO: Wrote 570000000 target and their corresponding decoy peptides
INFO: Wrote 580000000 target and their corresponding decoy peptides
INFO: Wrote 590000000 target and their corresponding decoy peptides
INFO: Wrote 600000000 target and their corresponding decoy peptides
INFO: Wrote 610000000 target and their corresponding decoy peptides
INFO: Wrote 620000000 target and their corresponding decoy peptides
INFO: Wrote 630000000 target and their corresponding decoy peptides
INFO: Wrote 640000000 target and their corresponding decoy peptides
INFO: Wrote 650000000 target and their corresponding decoy peptides
INFO: Wrote 660000000 target and their corresponding decoy peptides
INFO: Wrote 670000000 target and their corresponding decoy peptides
INFO: Wrote 680000000 target and their corresponding decoy peptides
INFO: Wrote 690000000 target and their corresponding decoy peptides
INFO: Wrote 700000000 target and their corresponding decoy peptides
INFO: Wrote 710000000 target and their corresponding decoy peptides
INFO: Wrote 720000000 target and their corresponding decoy peptides
INFO: Wrote 730000000 target and their corresponding decoy peptides
INFO: Wrote 740000000 target and their corresponding decoy peptides
INFO: Wrote 750000000 target and their corresponding decoy peptides
INFO: Wrote 760000000 target and their corresponding decoy peptides
INFO: Wrote 770000000 target and their corresponding decoy peptides
INFO: Wrote 780000000 target and their corresponding decoy peptides
INFO: Wrote 790000000 target and their corresponding decoy peptides
INFO: Wrote 800000000 target and their corresponding decoy peptides
INFO: Wrote 810000000 target and their corresponding decoy peptides
INFO: Wrote 820000000 target and their corresponding decoy peptides
INFO: Wrote 830000000 target and their corresponding decoy peptides
INFO: Wrote 840000000 target and their corresponding decoy peptides
INFO: Wrote 850000000 target and their corresponding decoy peptides
INFO: Wrote 860000000 target and their corresponding decoy peptides
INFO: Wrote 870000000 target and their corresponding decoy peptides
INFO: Wrote 880000000 target and their corresponding decoy peptides
INFO: Wrote 890000000 target and their corresponding decoy peptides
INFO: Wrote 900000000 target and their corresponding decoy peptides
INFO: Wrote 910000000 target and their corresponding decoy peptides
INFO: Wrote 920000000 target and their corresponding decoy peptides
INFO: Wrote 930000000 target and their corresponding decoy peptides
INFO: Wrote 940000000 target and their corresponding decoy peptides
INFO: Wrote 950000000 target and their corresponding decoy peptides
INFO: Wrote 960000000 target and their corresponding decoy peptides
INFO: Wrote 970000000 target and their corresponding decoy peptides
INFO: Wrote 980000000 target and their corresponding decoy peptides
INFO: Wrote 990000000 target and their corresponding decoy peptides
INFO: Wrote 1000000000 target and their corresponding decoy peptides
INFO: Wrote 1010000000 target and their corresponding decoy peptides
INFO: Wrote 1020000000 target and their corresponding decoy peptides
INFO: Wrote 1030000000 target and their corresponding decoy peptides
INFO: Wrote 1040000000 target and their corresponding decoy peptides
INFO: Wrote 1050000000 target and their corresponding decoy peptides
INFO: Wrote 1060000000 target and their corresponding decoy peptides
INFO: Wrote 1070000000 target and their corresponding decoy peptides
INFO: Wrote 1080000000 target and their corresponding decoy peptides
INFO: Wrote 1090000000 target and their corresponding decoy peptides
INFO: Wrote 1100000000 target and their corresponding decoy peptides
INFO: Wrote 1110000000 target and their corresponding decoy peptides
INFO: Wrote 1120000000 target and their corresponding decoy peptides
INFO: Wrote 1130000000 target and their corresponding decoy peptides
INFO: Wrote 1140000000 target and their corresponding decoy peptides
INFO: Wrote 1150000000 target and their corresponding decoy peptides
INFO: Wrote 1160000000 target and their corresponding decoy peptides
INFO: Wrote 1170000000 target and their corresponding decoy peptides
INFO: Failed to generate decoys for 61609 low complexity peptides.
INFO: Generated 1177005847 target peptides.
INFO: Generated 1176633139 decoy peptides.
INFO: Generated 2353638986 peptides in total.
INFO: Elapsed time: 9.26e+03 s
INFO: Finished crux tide-index.
INFO: Return Code:0
