## Supplementary figures and images for "Fast and memory efficient searching of large-scale mass spectrometry data using Tide"

### MSFragger_resource_usage.jpg

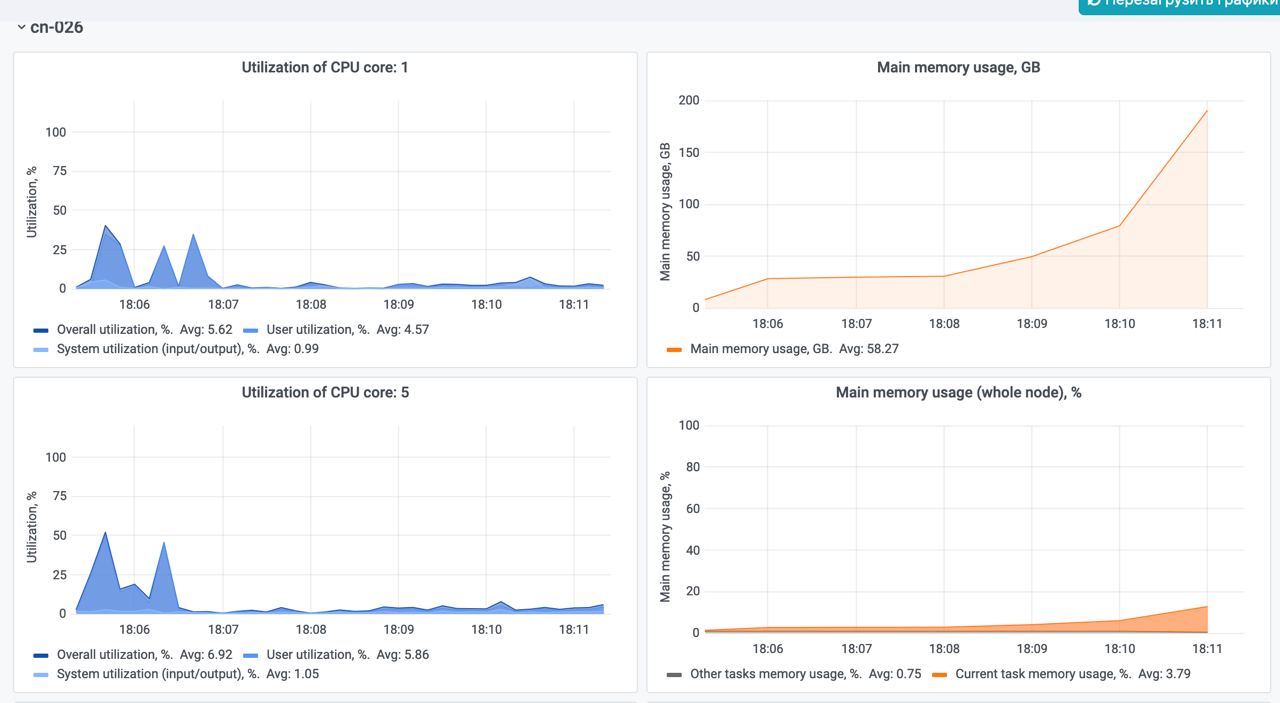

### SAGE_resource_usage.jpg

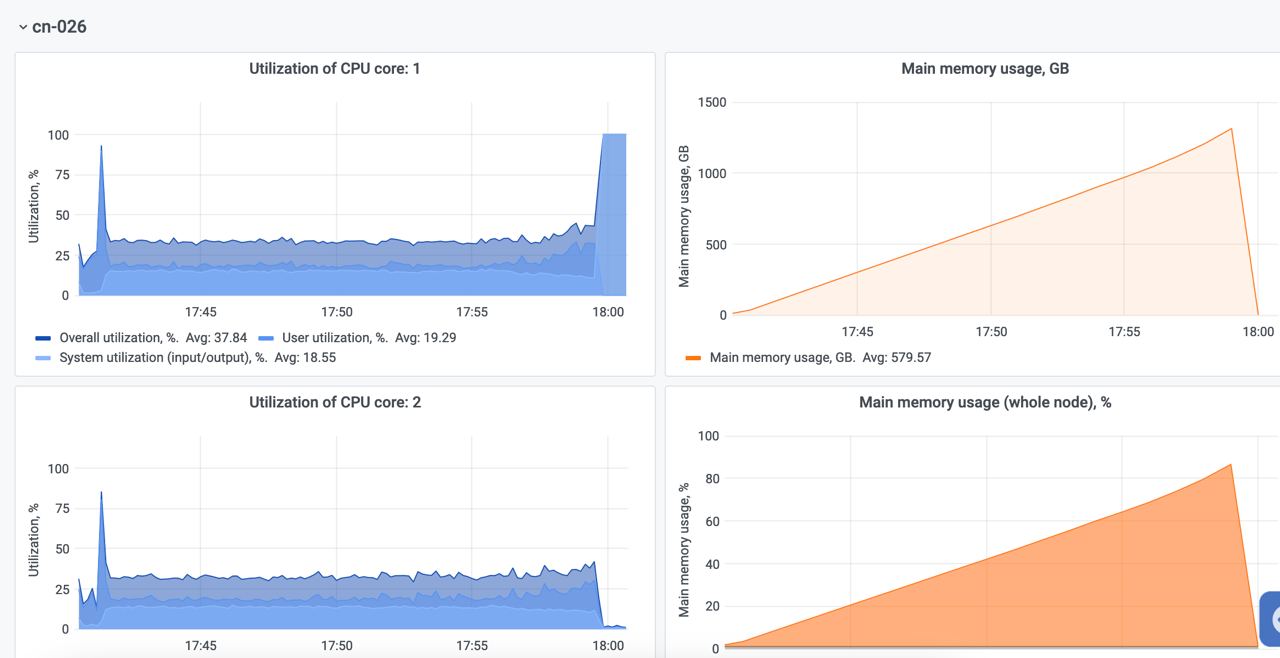
